# Exceptional Late Devonian arthropods document the origin of decapod crustaceans

**DOI:** 10.1101/2020.10.23.352971

**Authors:** Pierre Gueriau, Štěpán Rak, Krzysztof Broda, Tomáš Kumpan, Tomáš Viktorýn, Petr Valach, Michał Zatoń, Sylvain Charbonnier, Javier Luque

## Abstract

With over 15,000 extant species, Decapoda—or ten-legged crustaceans such as crabs, shrimp, lobsters, and relatives— are among the most speciose and economically important group of crustaceans. Despite of their diversity, anatomical disparity, and remarkable fossil record extending back to the Late Paleozoic, the origins of Decapoda and their phylogenetic relationships with eumalacostracans remains elusive and inconclusive. Molecular dating suggests that decapods originated in the Late Ordovician (~450 Mya), but no reliable fossil crown groups are found until the Late Devonian. Moreover, there is no consensus on which lineages belong to stem groups, obscuring our understanding of the roots of the ten-legged decapod body plans as a whole, and how they relate to other non-decapod crustaceans. We present new, exceptional fossils from the Late Devonian of Czech Republic and Poland that belong to †Angustidontida, an odd shrimp-looking crustacean with a combination of anatomical features unlike those of any crown eumalacostracan known—extinct or extant. Our phylogenetic analyses, including representatives of all major lineages of crown eumalacostracans plus †Angustidontida, identify angustidontids as the only known stem-group decapod, and give hints about the transformation series, polarity of change, and evolutionary pathways leading to the modern decapod body plans seen today.

## 1. Introduction

Among crustaceans, the representatives of the order Decapoda are easily recognizable due to their distinctive body forms. They have a carapace that is fused to the underlying thoracic segments, an abdomen (pleon) constituted by six segments or pleonites, and most lineages other than crabs have a telson and uropods that often form a tailfan. Moreover, crown-group decapods have eight pairs of thoracic appendages or thoracopods, from which the first three pairs are modified into maxillipeds for feeding, while the remaining five pairs are developed into true thoracic legs (pereopods), hence the name of the order (Decapoda = ten-footed crustaceans) (e.g. [1]). Despite of such a distinctive body arrangement, the deep relationships among decapods have long been conflicting and unresolved. Recent phylogenomic analyses with assembled sequence data from 94 species (including 11 of the 12 major lineages) recovered a robust extant decapod tree of life [2]. Unfortunately, the origin of Decapoda remains obscure as i) there is no consensus on the sister group to crown-group Decapoda, and ii) no stem-group decapod forms have been confirmed from reliable Paleozoic (Late Devonian-Carboniferous) fossils [3], with the latter being too derived or insufficiently preserved to be informative about the origins of the total-group Decapoda [1,3,4]. Furthermore, molecular divergence times suggest that crown decapods diverged in the Late Ordovician, leaving a gap of nearly 70 million years between their estimated split and their earliest confirmed fossils, and implying a significant cryptic Paleozoic history [2].

Among the few known middle Paleozoic eumalacostracan crustaceans, the iconic shrimp-looking †Angustidontida from Late Devonian and Early Carboniferous of North America and Europe have been envisioned as closely related to decapods [1,2,4,5]. Unlike crown decapods, †Angustidontida bears one or two pairs of elongated comb-like maxillipeds (figure 1*a*; one pair in †*Angustidontus* Cooper, 1936, two pairs in †*Schramidontus* Gueriau, Charbonnier & Clément, 2014 [4]). Isolated remains of such comb-like maxilliped were described as either actinopterygian fish jaws, eurypterid raptorial appendages, or crustacean appendages [1, 5], until more complete and articulated material confirmed their crustacean affinities. †Angustidontida is morphologically similar to Decapoda [4], but differs from it in the number of maxillipeds and the size and connection of the pleonal segments (uniform in angustidontids; variable in most decapods, with an expanded second pleonal pleuron and an enlarged third pleonal somite). In particular, angustidontids exhibit remarkable similarity with †*Palaeopalaemon newberryi* Whitfield, 1880, another fossil crustacean from the Late Devonian recognized undoubtedly as a crown decapod [6,7], yet †*P. newberryi* is distinct from angustidontids [7]. Gueriau et al. [4] postulated that angustidontids were early decapods that filled the gap between krill or Euphausiacea (no maxillipeds) and Decapoda (three pairs of maxillipeds), and that they were closer to Decapoda than to the extant *Amphionides reynaudii* Milne Edwards, 1832 (a rare crustacean bearing a single pair of maxillipeds). This apomorphic trait placed *Amphionides* as the sister taxon to Decapoda, and it was so unique that merited its own order Amphionidacea [8]. The single pair of maxillipeds of *A. reynaudii* told a beautiful story about the possible origins of Decapoda, and filled the gap between Euphausiacea and Decapoda, but it took an unexpected turn when molecular phylogenetics identified Amphionidacea as the larval form of a caridean decapod shrimp [9]. As a consequence, †Angustidontida stood as the only remaining sister group candidate to Decapoda, or at least the only extinct lineage recognizable between euphausiaceans and decapods [1]. However, the angustidontid material described so far does not exhibit any character autapomorphic of Decapoda [4], for which new material and a revision in a more explicit phylogenetic framework is still lacking [2].

**Figure 1.**
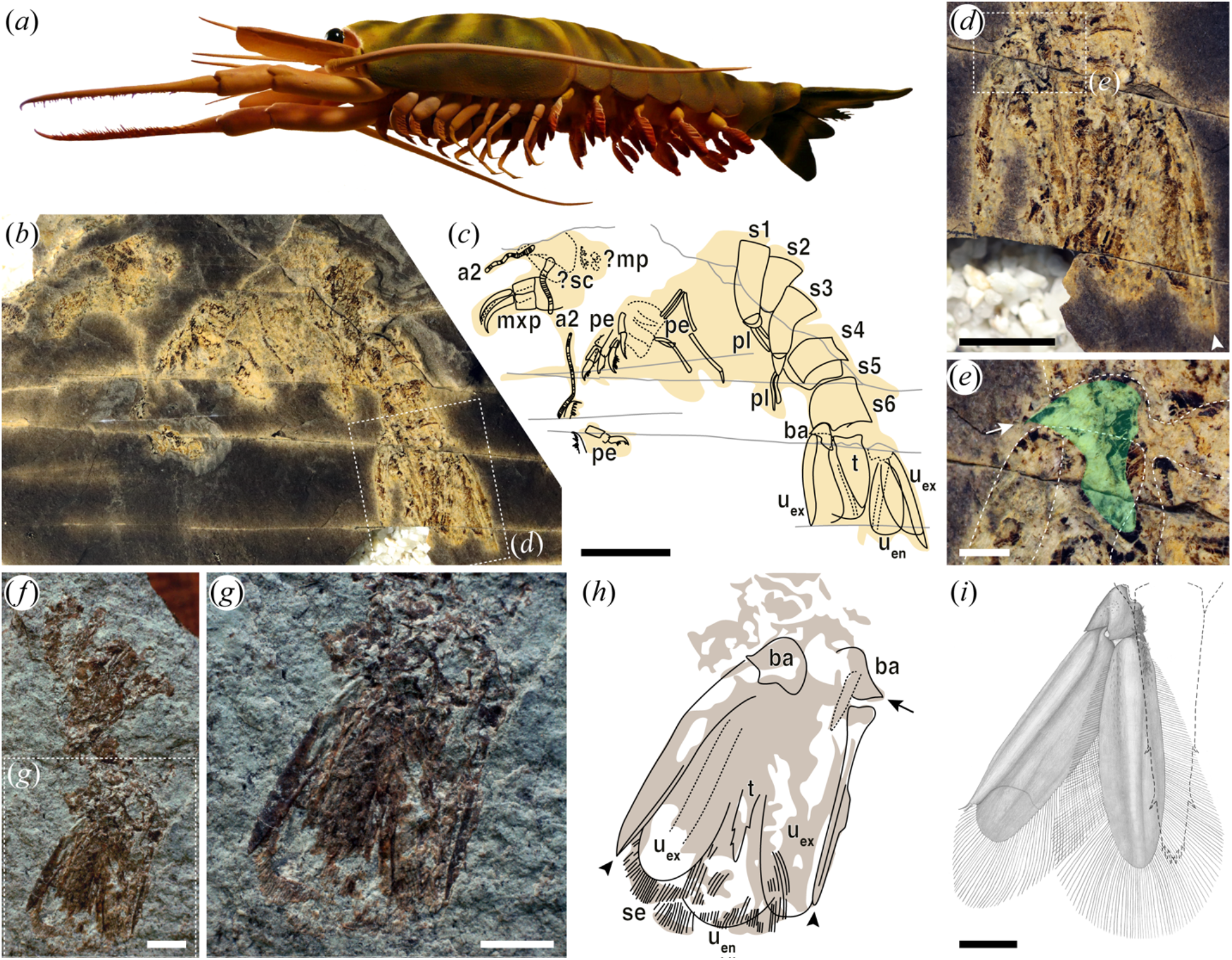
Angustidontid tailfan anatomy. (*a*) Reconstruction of †*Angustidontus* in ventro-lateral view; SHAPE-PLAST thermoplastic model by T.V. (*b–e*) †*Angustidontus* aff. *moravicus*, H33/1A, complete specimen (likely an exuvia) in dorso-lateral view from the Hády quarry (lower Fammenian, Czech Republic). (*b*) Photo of specimen covered in water. (*c*) Interpretative line drawing. (*d*) Close-up of the tailfan, from the box area in *b*. (*e*) Close-up of the basipod (highlighted in green), from the box area in *d*. (*f–h*) †*Angustidontus* aff. *seriatus*, GIUS 4-3622/kor112, articulated pleon in dorso-lateral view from the Kowala quarry (lower Fammenian, Holy Cross Mountains, Poland). (*f*) Photo dry. (*g*) Close-up of the tailfain, from the box area in *f*. (*h*) Interpretative line drawing. (*i*) Pencil drawing of the uropods of the extant shrimp *Crangon crangon* Linnaeus, 1758. Illustration by Dr. Verena Kutschera. Abbreviations: a2, antenna; ba, basipod; ?mp, ?mouth parts; mxp, maxillipeds; pe, pereopods; pl, pleopods; s1-s6; pleonal somites 1 to 6; se, setae; ?sc, ?scaphocerites; t, telson; u_en_, uropodal endopod; u_ex_, uropodal exopod. Arrowheads and arrows point out the latero-distal spiky outgrowth of u_ex_ carina, and the latero-distal tipped prolongation of the basipod, respectively. Scale bars, 1 cm in *b* and *c*, 5 mm in *d*, 2 mm in *f–h*, and 1 mm in *e* and *i*.

Here, we describe new anatomical features of †Angustidontida based on new, exceptional fossils from the Late Devonian of Czech Republic and Poland, showing in exceptional detail pleonal and caudal features that allow us to explore their phylogenetic position within Malacostraca, and its systematic implications for understanding the origins of crown decapod crustaceans.

## 2. Materials and methods

### (a) Material

The fossil specimens investigated herein come from the Hády quarry (lower Fammenian, Czech Republic; see next section for geological settings) and the Kowala quarry (lower Fammenian, Holy Cross Mountains, Poland; see [10] for geological settings), and are housed at the Czech Geological Survey, Prague, and the Institute of Earth Sciences, University of Silesia, Sosnowiec, respectively. Fossils from the Hády quarry presented herein (figure 1*b*–*e* and electronic supplementary material, figure S1) have never been published or mentioned before; the specimen from the Kowala quarry (figure 1*f*–*h*) have been mentioned, but not figured in [10]. They were studied under a binocular microscope, both dry and covered in water with a low angle light to better reveal relief, and photographed with a SLR camera coupled with macro lens equipped with polarizing filters. Interpretative line drawings were produced on the photographs while observing the specimens with different light angles under the binocular microscope. Drawings and figures were made using Adobe Illustrator.

### (b) Geological settings and stratigraphy of the Hády quarry, Czech Republic

The new angustidontid material from Czech Republic comes from section H33 of the Hády quarry (part Městký lom), northeastern outskirts of Brno, Czech Republic. The Hády quarry exposes a southernmost outcrop of the Moravian Karst Paleozoic, belonging to the Variscan Brunovistulian Unit. Devonian and Carboniferous sedimentary rocks of the Brunovistulian Unit were deposited at a southern margin of Laurussia [11,12,13,14].

In the Hády quarry, Late Frasnian pure massive limestones of the Macocha Formation are overlain by the latest Frasnian to late Famennian nodular to well bedded limestones of the Líšeň Formation (electronic supplementary material, figure S1). A drowning of the Frasnian outer ramp and transition to an Famennian hemipelagic slope environment with turbiditic influence is recorded in the studied section [15,16,17]). Angustidontid-bearing bed is one of the calcareous shale intercalations, which alternate with platy calcilutites and occasional banks of medium-grained calcirudites (electronic supplementary material, figure S1). The calcareous shales represent “background” hemipelagic facies deposited in relatively low-energy slope environment, well below storm-wave base. The platy limestones were interpreted as calciturbidites (e.g. [16,18,19]). Limestone sample just below the angustidontid-bearing shale yielded 110 conodonts, which are typical for the lower Famennian *Palmatolepis minuta minuta* to *Palmatolepis crepida* conodont zones interval (*sensu* [20]), involving *Pa. triangularis*, *Pa. minuta minuta, Pa. delicatula delicatula* and *Icriodus alternatus*.

### (c) Phylogenetic analysis

We investigated the phylogenetic position of †Angustidontida using a modified morphological dataset for malacostracan crustaceans after [8] (characters 1–93). Following [21], we added four additional characters related to the morphology of the basipods and uropods (characters 94–97). The operational taxonomic unit (OTU) ‘Amphionidacea’, included in the analyses of [8], was culled from our analysis as it has been demonstrated that amphionidaceans are the larvae of a decapod shrimp rather than a distinct malacostracan order [1, 9]. The final data matrix, containing 19 OTUs (3 in outgroup, 16 in ingroup) and 97 adult morphological characters, was built in Mesquite 3.51 [22]. Undetermined or not preserved characters were scored as ‘?’, and inapplicable characters as ‘–‘. Multiple character states present in a given OTU were scored as polymorphisms.

We analyzed the dataset using Bayesian inference (BI) as implemented in MrBayes v.3.2.5 [23]. The dataset was analyzed under the traditional Mk model [24] with an ascertainment bias correction to account for scoring only variable morphological characters, and gamma distributed rate variation. Each analysis was performed with two independent runs of 3×10^7^ generations each. We used the default settings of four chains per independent run. The relative burn-in fraction was set to 25%, and the chains were sampled every 200 generations. We used Tracer v. 1.7.1 [25] to determine whether the runs reached stationary phase and to ensure that the effective sample size for each parameter was greater than 200. Results of the Bayesian runs were summarized as a majority-rule consensus tree of the post-burnin sample (figure 2).

**Figure 2.**
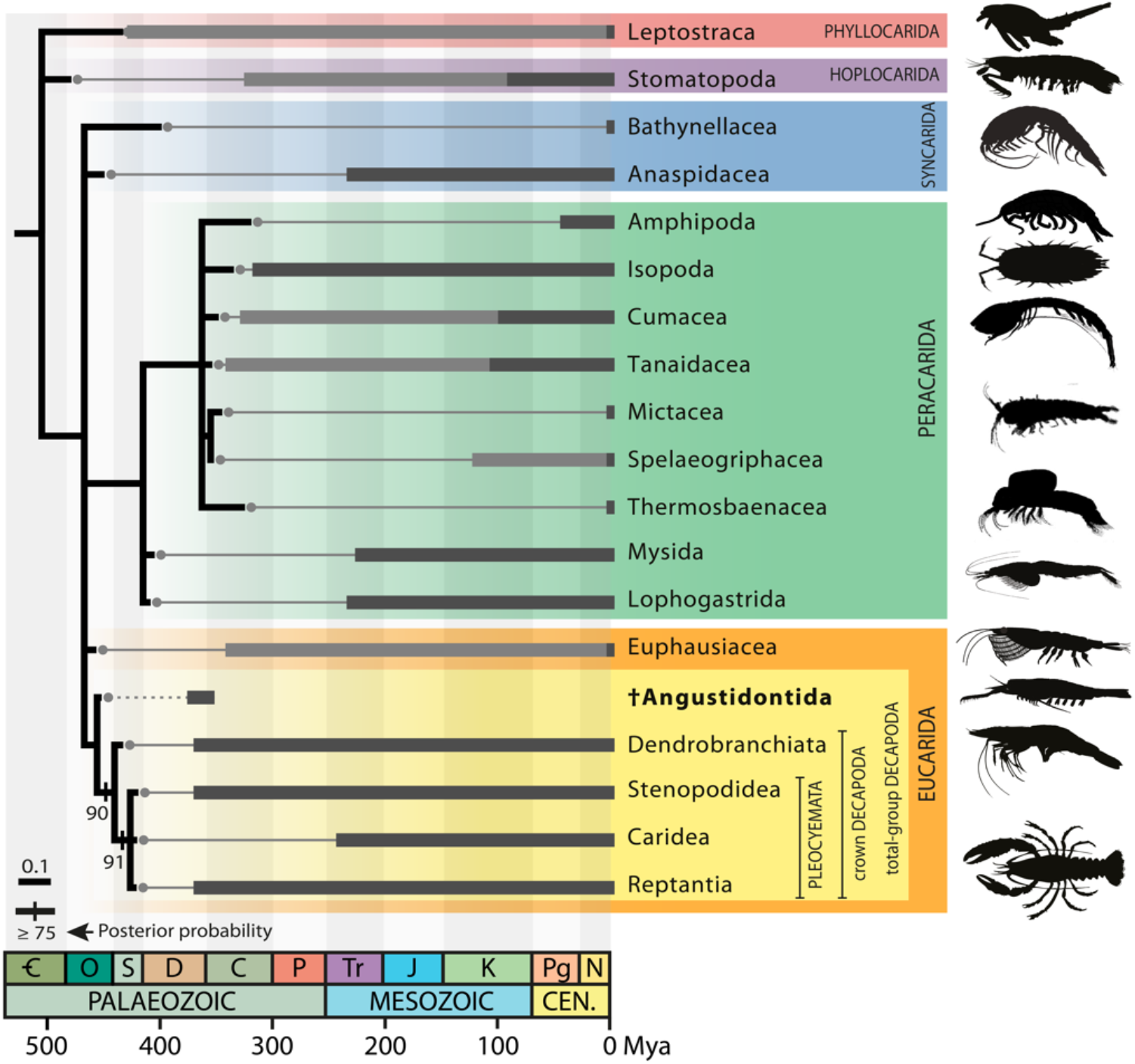
Phylogenetic position of angustidontids within malacostracan crustaceans. Bayesian majority-rule consensus topology of the post-burnin sample of trees, plotted on stratigraphy. The obtained tree (black line) was scaled to accommodate a Late Ordovician divergence date for Decapoda [2] and the stratigraphic ranges of terminal taxa (light and dark grey thick lines for stem- and crown-groups, respectively; see electronic supplementary material, table S1, for ages justification) here constrained by the first occurrence of stem Leptostraca. Posterior probability support values indicated above branches. Branches with posterior probability support < 75% are collapsed.

We also conducted maximum parsimony (MP) phylogenetic analyses in TNT v.1.5 [26]. The data was analyzed under implicit enumeration for equally weighted maximum parsimony (EWMP) (figure S2*a*), and different implied weights (K=3, 6, 12) as additional tests of placement of †Angustidontida (electronic supplementary material, figure S2*b–d*). Bootstrap and jackknife values were calculated after 10,000 replications each using default settings. Bremer support values for the EWMP implicit enumeration were calculated under tree bisection reconnection, and retained trees suboptimal by 30 steps. All characters were unordered.

## 3. Results and discussion

New material assigned to †*Angustidontus* aff. *moravicus* Chlupáč, 1978 [11] collected from Czech Republic (figure 1*b*–*e* and electronic supplementary material, figure S1), and †*Angustidontus* aff. *seriatus* Cooper, 1936 from Poland (figure 1*f*–*h*) reveals fine details of the anatomy of their tailfan previously unknown, which in turn allow for a detailed taxonomic and phylogenetic comparison with other malacostracans. The angustidontid tailfan consists of a triangular telson flanked by two pairs of leaf-shaped uropods (figure 1*b*–*d*,*g*,*h*; see also [4,5]). The uropodal exopod bears a carina, and prolongs latero-distally in a non-articulated, spiky outgrowth (figure 1*d*,*h* arrowheads). The lateral, distal, and median margins of the endopod and exopod (up to the spiky outgrowth for the latter) are surrounded by plumose setae of different length, longest distally and shortest proximo-medially and latero-distally (figure 1*g*,*h*). The basipod is sub-conical and possesses a latero-distal, tipped prolongation (figure 1*e*,*h* arrows; see also [5]). According to Kutschera et al. [21]’s work on the phylogenetic signal conveyed by malacostracan basipods and uropods morphology, such combination of characters is unique to Decapoda. It is worth noting that, as described by Rolfe and Dzik [5], the uropodal exopod is not divided into two portions by a medio-lateral suture or diaeresis, a feature also absent in Euphausiacea and Dendrobranchiata (though some fossils assigned to the group do possess one [27,28,29]) but found in Caridea [21]. The presence or absence of a longitudinal median keel on the basipod, specific to Decapoda [21], is impossible to determine on our flattened fossil material. Aside this, the anatomy of the angustidontid tailfan is virtually identical to that of decapods (figure 1*i*), adding support to an alliance between these groups.

Bayesian inference and maximum parsimony phylogenetic analyses recover †Angustidontida as the sister group to the crown-group Decapoda (figure 2, and electronic supplementary material, figure S2). Besides filling the gap between Euphausiacea and Decapoda, an affiliation of angustidontids as the closest known sister taxon to crown group Decapoda, i.e. as part of total-group Decapoda, has important implications for reconciling molecular divergence dates for decapods (Late Ordovician [2]) and their earliest known fossil record (Late Devonian [1,3,4,6,7]). Our findings identify angustidontids as the only stem-group decapod known to date, and suggest that their apparently cryptic Paleozoic history [2] might be an artifact of the overlooked disparate stem-decapod body plans, compared to crown-group forms. Our results indicate that the origin of decapod-like eucaridans lies within more than ten-footed crustaceans, and gives hints about the transformation series, polarity of change, and evolutionary pathways leading to the characteristic decapod body arrangement and thoracopod configuration seen across the main crown groups as recognized today.

## Supporting information

Electronic supplementary material

## Additional information

### Data accessibility

Requests for materials should be addressed to Petr Budil (petr.budil@geologycz) or M.Z. for the Hády quarry and the Kowala quarry, respectively. The phylogenetic data matrix is available in Morphobank http://morphobank.org/permalink/?P3739. Additional data are available as electronic supplementary material.

### Authors’ contributions

P.G. and S.R. conceived the project. S.R., K.B., T.K., T.V. and P.V. collected the material from Czech Republic and composed the geological settings and Figure S1A. M.Z. collected and provided photographs of the specimen from Poland. P.G., S.C. and J.L. defined the phylogenetic characters and coded the matrix. J.L. conducted the phylogenetic analyses. P.G., S.R. and J.L. discussed the results and P.G. prepared the manuscript with input from all other authors.

### Competing interests

The authors declare no competing interests.

### Funding

Part of this study was supported by the research project 19-17435S of the Czech Science Foundation (GAČR). K.B. was supported by the National Science Center grant PRELUDIUM (2015/19/N/ST10/01527). J.L. thanks the Natural Science and Engineering Research Council of Canada Postdoctoral Fellowship (NSERC PDF), the Yale Institute for Biospheric Studies (YIBS), and the National Science Foundation Grant DEB #1856679 (USA).

## Acknowledgements

We thank Dr. Verena Kutschera for the pencil drawing of the uropods of *Crangon crangon* presented in figure 1*i* and Dr. Frederick R. Schram for comments and suggestions about the phylogenetic analysis at an early stage of this research.

## Notes

http://morphobank.org/permalink/?P3739

### Competing Interest Statement

The authors have declared no competing interest.

