## Supplementary material for "Exceptional Late Devonian arthropods document the origin of decapod crustaceans": Electronic supplementary material

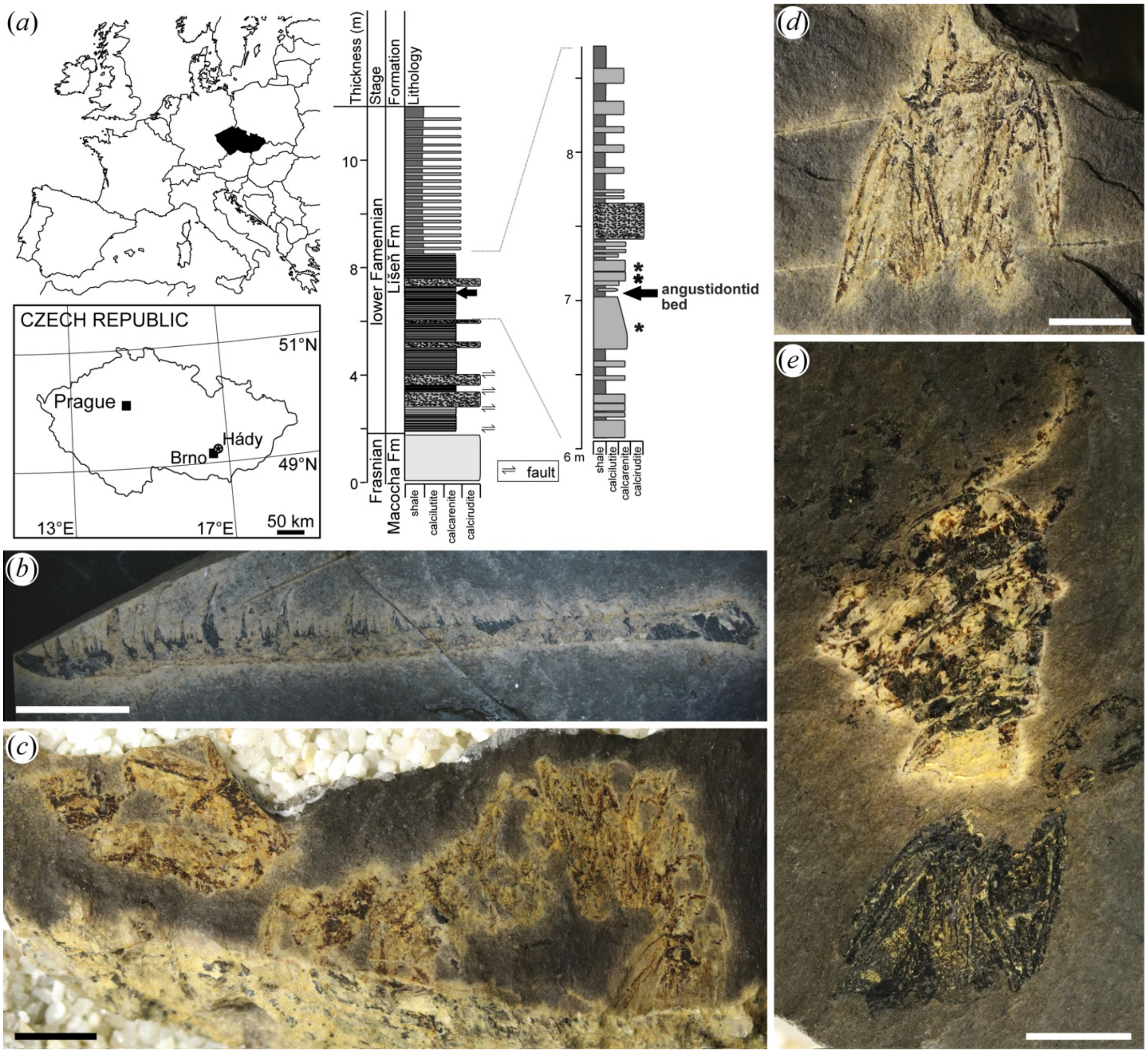


**Figure S1.** Stratigraphy of the Hády quarry (lower Fammenian, Czech Republic) and additional angustidontid material. (*a*) Location and lithological and stratigraphical log of the Hády quarry H33 section yielding the angustidontid bed. Asterisks show locations of conodont samples (lower Famennian *Palmatolepis minuta minuta to Palmatolepis crepida* conodont zones interval). (*b*–*e*) Additional specimens of *Angustidontus* aff. *moravicus* Chlupáč, 1978. (*b*) H33/60A, isolated maxilliped. Photo dry. (*c*) H33/160B, articulated exuvia in dorso-lateral view. Photo covered in water. (*d*) H33/1C, counterpart of the tailfan shown in figure 1*b*–*e*. Photo dry. (*e*) H33/2A, articulated pleon and tailfan in dorsal view. Photo covered in water. Scale bars, 5 mm.

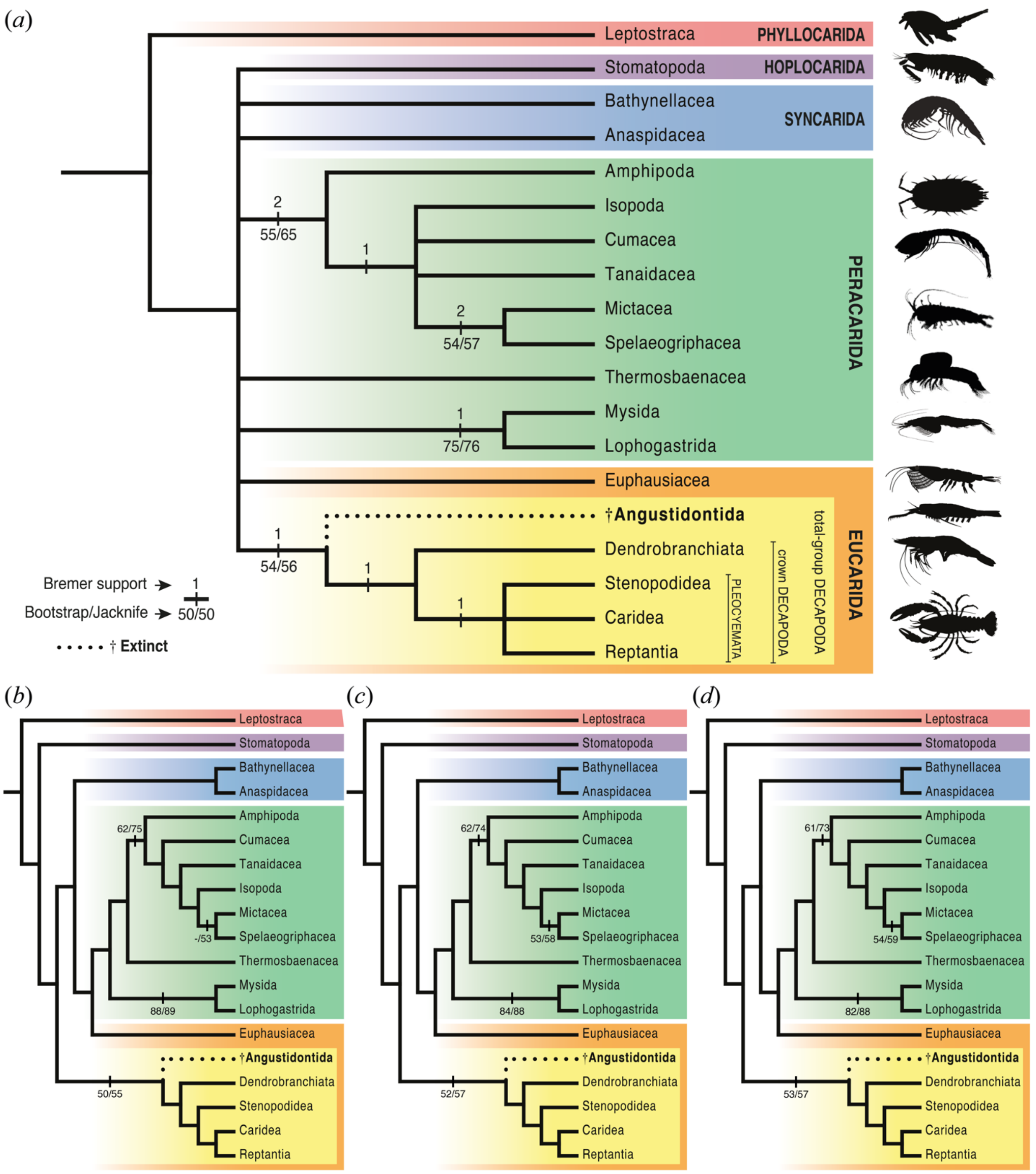


**Figure S2.** Maximum parsimony placement of †Angustidontida within malacostracan crustaceans. (*a*) Equally-weighted parsimony (EWMP): strict consensus of five most parsimonious trees. (*b*–*d*) Implied-weights maximum parsimony (IWMP) topologies with concavity values of K=3 (*b*) (fit= 21.793), K=6 (*c*) (fit= 13.393), and K=12 (*d*) (fit= 7.592). Analyses performed in TNT v.1.5.

**Table S1.** First appearance datum (FAD) ages justification for the stratigraphic ranges of the stem- and crown-group malacostracan operational taxonomic units (OTUs) used in the phylogenetic analyses.

| OTU | | FAD taxon | Stage / Age | Minimum absolute age (Ma) | Reference(s) |
|---|---|---|---|---|---|
| Leptostraca | stem | *Cinerocaris magnifica*; *Cascolus ravitis* | Sheinwoodian-Homerian | 429.8 | a, b, [1] |
| | crown | | – | | [1] |
| Stomatopoda | stem | *Daidal acanthocercus* | Middle Pennsylvanian | 306.9 | c, d |
| | crown | *Ursquilla yehoachi* | uppermost early Campanian | 79.01 | a |
| Bathynellacea | stem | | – | | e |
| | crown | | – | | e |
| Anaspidacea | stem | | no evidence for older stem forms | | [1] |
| | crown | *Anaspidites antiquus* | Anisian | 240.5 | a, [1] |
| Amphipoda | stem | | no evidence for older stem forms | | [1] |
| | crown | *Gammaroidorum vonki* | upper Hauterivian | 129.4 | f |
| Isopoda | stem | | no evidence for older stem forms | | [1] |
| | crown | *Hesslerella shermani* | uppermost Moscovian | 306.9 | a |
| Cumacea | stem | *Securicaris spinosus*; *Carbocuma imoensis*; *Ophthalmdiastylis parvulorostrum* | Upper Mississipian | 322.8 | g |
| | crown | *Eobodotria muisca* | upper Cenomanian | 93.7 | g |
| Tanaidacea | stem | *Anthracocaris scotica* | Visean | 330.7 | [1] |
| | crown | *Alavatanais carabe* | upper Albian | 100.1 | a |
| Mictacea | stem | | – | | [1] |
| | crown | | – | | [1] |
| Spelaeogriphacea | stem | *Spinogriphus ibericus* | upper Barremian | 125 | h |
| | crown | | – | | h |
| Thermosbaenacea | stem | | – | | i |
| | crown | | – | | i |

| | | | | | |
|---|---|---|---|---|---|
| Mysida | stem | | no evidence for older stem forms | | |
| | crown | *Aviamysis pinetellensis* | upper Ladinian | 235.0 | j |
| Lophogastrida | stem | | no evidence for older stem forms | | |
| | crown | *Yunnanocopia grandis*; *Y. longicauda* | Anisian | 240.5 | k |
| Euphausiacea | stem | *Anthracophausia dunsiana* | Visean | 330.7 | [1] |
| | crown | | – | | l |
| †Angustidontida | | *Angustidontus seriatus* | upper Frasnian–Lower Mississippian* | 370.6–358.5* | m, n |
| Dendrobranchiata | stem | | no evidence for older stem forms | | |
| | crown | *Aciculopoda mapesi* | Fammenian | 358.5 | o, [1] |
| Stenopodidea | stem | | no evidence for older stem forms | | |
| | crown | *Devonostenopus pennsylvanensis* | Upper Devonian | 358.5 | p, [1] |
| Caridea | stem | | no evidence for older stem forms | | |
| | crown | formally undescribed | middle Olenekian | 247.2 | q |
| Reptantia | stem | | no evidence for older stem forms | | |
| | crown | *Palaeopalaemon newberry* | lupper Fammenian | 358.5 | a, [1] |

* First and last occurrences of †Angustidontida, the only extinct order studied herein.
